# Fast and accurate joint inference of coancestry parameters for populations and/or individuals

**DOI:** 10.1101/2022.01.28.478138

**Authors:** Tristan Mary-Huard, David Balding

## Abstract

We introduce a fast, new algorithm for inferring jointly the *F*_*ST*_ parameters describing genetic distances among a set of populations and/or unrelated diploid individuals, and a tree representing their genetic structure, from allele count data. While the inferred tree typically reflects historical processes of splitting and divergence, its aim is to represent the actual genetic variance, with *F*_*ST*_ values specified by branch lengths. We generalise two major approaches to defining *F*_*ST*_, via correlations and mismatch probabilities of sampled allele pairs, which measure shared and non-shared components of genetic variance. A diploid individual can be treated as a population of two gametes, which allows inference of coancestry coefficients for individuals as well as for populations, or a combination of the two. A simulation study illustrates that our fast method-of-moments estimation of *F*_*ST*_ values, simultaneously for multiple populations/individuals, gains statistical efficiency over pairwise approaches by pooling information about ancestral allele frequencies. We apply our approach to genome-wide genotypes from the 26 worldwide human populations of the 1000 Genomes Project. We first analyse at the population level, then a subset of individuals and in a final analysis we pool individuals from the more homogeneous populations. This flexible analysis approach gives many advantages over traditional approaches to population structure/coancestry, including visual and quantitative assessments of long-standing questions about the relative magnitudes of within- and between-population genetic differences.

**Author summary:** We propose new ways to measure, and visualise in a tree, the genetic distances among a set of populations using allele frequency data. The two genomes within a diploid individual can be treated as a small population, which allows a flexible framework for investigating genetic variation within and between populations. Genetic structure can be accurately and efficiently represented in a tree with nodes representing either homogeneous populations or genetically diverse individuals, for example due to admixture. We first generalise the long-established measure of genetic distance, *F*_*ST*_, to tree-structured populations and individuals, finding that two measures are required for each pair of populations, corresponding to their shared and and non-shared genetic variation. We show using a simulation study that our novel tree-based estimators are more efficient than current pairwise estimators, and we illustrate the potential for novel ways to explore and visualise genetic variation within and between populations using a worldwide human genetic dataset.

## Introduction

*F*_*ST*_ is a measure of between-population genetic distance introduced in the seminal work of [1]. Several definitions have been proposed, for example in terms of correlations of alleles sampled in observed populations, relative to an actual or hypothetical reference population, or in terms of average mismatch probabilities for pairs of alleles from the same population, and from different populations. Different underlying definitions have complicated comparisons of the many *F*_*ST*_ estimators that have been proposed. These include sum-of-squares estimators in a components-of-variance framework [2, 3], and maximum likelihood estimation based on the variance parameter of the multinomial-Dirichlet distribution (beta-binomial for diallelic markers) [4]. Bhatia *et al*. [5] used a moment estimator of mismatch probabilities, similar to the estimators of Hudson [6] and Nei [7].

With very large numbers of single-nucleotide variants (SNV), estimators can be precise and so differences in definitions can be important. *F*_*ST*_ estimates for pairs of worldwide human populations have differed by almost a factor of two even when based on the same dataset, due to sensitivity to the minor allele fractions (MAF) [5]. The estimator of [5] is simple and computationally fast, and helped to resolve the controversy over estimation of *F*_*ST*_ for pairs of populations given genome-wide SNV data.

Currently, researchers with multi-population data typically apply a standard estimator separately for each pair of populations. Recent advances [8, 9], following earlier suggestions [10, 11], have added flexibility through integrating the analyses of individuals and populations. Here we propose fast and statistically-efficient method-of-moments estimation of *F*_*ST*_, simultaneously for multiple populations and/or individuals, by inferring a tree of ancestral populations that models shared and unshared components of allele-frequency variances via tree branch lengths that specify *F*_*ST*_ values.

Unrelated diploid individuals can be treated as populations of two gametes. While accurate tree inference with all leaves corresponding to individuals is infeasible for large sample sizes, a hybrid approach can be used with some leaves corresponding to homogeneous populations and others to individuals with greater genetic diversity, perhaps due to admixture. A flexible modelling framework using a sequence of analyses can be employed to converge on a best-fitting representation of population structure. Here, we do not model the effects of linkage, and so we only consider individuals with no very recent shared ancestors. To introduce our new approach as simply as possible, we do not model inbreeding here.

While the tree typically reflects evolutionary history, it primarily provides a visual representation of the actual genetic variance inferred from observed allele frequencies. Many authors interpret coancestry parameters in terms of identity-by-descent (IBD) probabilities [9, 12, 13]. The IBD concept is popular and allows an intuitive language, but can be problematic [14] because there is a common ancestor at each genome site and different approaches are used to convert the continuous time-since-common-ancestor into a binary IBD state. These include current or ancestral reference populations [8, 9, 13] or mutation events [11, 15]. Our coancestry parameters describe components of allele frequency variance, and we use IBD terminology to assist with interpretation and for compatability with other approaches. Our framework is similar to the IBD-centred approach of [9] in that reference allele frequencies are assumed to be those of the most recent population ancestral to the sampled populations. It follows that *F*_*ST*_ ≥ 0 here, as in [9] but in contrast with other approaches that allow *F*_*ST*_ < 0 [5, 8].

We first generalise to multiple populations the correlation and mismatch definitions, denoted 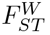 and 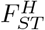, respectively, the superscripts referring to the seminal authors Weir/Wright and Hudson. For tree-structured populations, 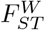 and 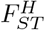 capture complementary aspects of population structure, corresponding to the lengths of shared and non-shared branches between the observed populations and the inferred ancestral population. Our inferred trees are binary, but since zero branch lengths are allowed, more general tree structures are possible.

We illustrate our approach via inference of *F*_*ST*_ in simulated populations, finding that our novel tree-based estimator of 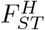 outperforms the pairwise estimator [5] in most comparisons, because it draws information from all observed populations, not only the two being compared. This advantage is greatest in the lowest-information setting that we consider with only 66 SNVs and 10 gametes (haploid individuals) sampled per population. We also analyse the 1000 Genomes Project data and derive a tree-based representation of the genetic variation among the 26 populations that reveals important insights. We further investigate 6 of these populations using individual coancestry coefficients, contrasting visually and quantitatively the within- and between-population genetic differences.

## Materials and methods

### Statistical model and definitions of *F*_*ST*_ in the classical setting

Assuming the independent-descent population model (Fig. 1), write *f* for the (unknown) reference allele fraction in Population 0 at a given locus, while *f*_*k*_ denotes its value in Population *k*. We assume 𝔼 [*f*_*k*_|*f*] = *f* and:

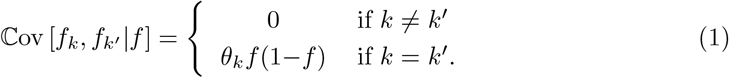

where the parameter *θ*_*k*_ ∈ [0, 1], closely related to *F*_*ST*_ (see below), measures the divergence of Population *k* from Population 0 due to drift, mutation, migration and selection. Ignoring mutation, *θ*_*k*_ can be interpreted as the probability that the sampled alleles are IBD from a common ancestor between Populations 0 and *k*.

**Figure 1.**
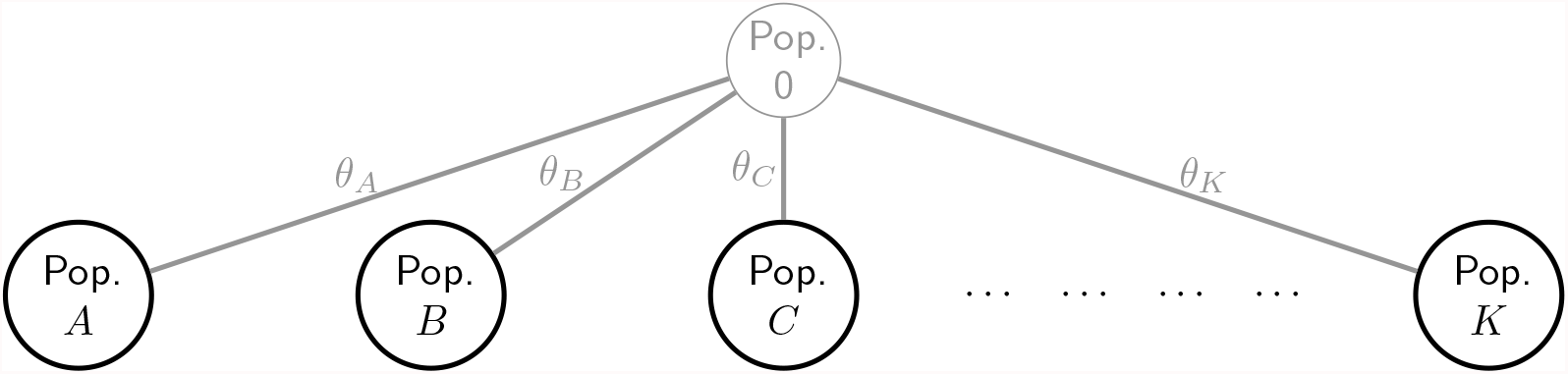
The independent-descent population model. [2, 3]. The ancestral population (Population 0) is unobserved. Allele count data are available from each of the other populations, which are assumed to have evolved independently from Population 0, with the level of divergence reflected in the *θ* values.

Let *x*_*k*_, *y*_*k*′_ ∈ {0, 1} be indicators of the reference allele for random allele draws at the locus in populations *k* and *k*^′^. We assume 𝔼 [*x*_*k*_|*f*_*k*_] = *f*_*k*_, 𝔼 [*y*_*k*′_ |*f*_*k*′_] = *f*_*k*′_ and ℂov [*x*_*k*_, *y*_*k*′_ |*f*_*k*_, *f*_*k*′_] = 0. Following Weir-Hill [3], and close to Wright [1], we define:

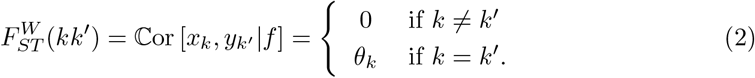

In the case *k* = *k*^′^, we will write *k* in place of *kk*^′^. Correlations can in general be negative, and *F*_*ST*_ < 0 could arise if the reference population were not ancestral to those sampled [8]. We also define the population analogue of the Hudson estimator [5, 6] as

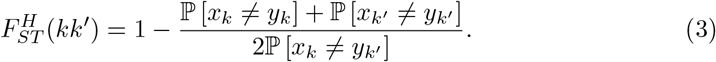

for *k* ≈ *k*′, with 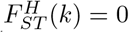.

Whereas 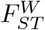 is a correlation of sampled alleles, 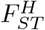 is based on mismatch probabilities, or expected heterozygosity, within and between two sampled populations. Under the independent-descent model, (1) leads to ℙ[*x*_*k*_ ≠*y*_*k*′_] = *f* (1−*f*) and ℙ[*x*_*k*_ ≠ *y*_*k*_] = (1−*θ*_*k*_)*f* (1−*f*), so that

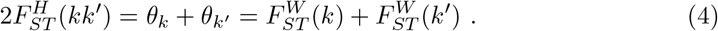

While 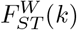 measures the divergence of Population *k* from Population 0, 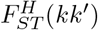 is the average divergence of Populations *k* and *k*^′^. We will see below that in more complex multi-population settings 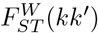 and 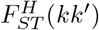 measure, respectively shared and non-shared genetic variation in populations *k* and *k*^′^.

### Generalisation of 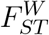 and 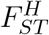 to tree-structured populations

First we assume a pre-specified tree topology (see Fig. 2 for an example), we express 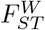 and 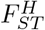 in terms of the *θ* parameters and propose estimators. Then we develop a method for joint inference of the tree topology and the *θ*.

**Figure 2.**
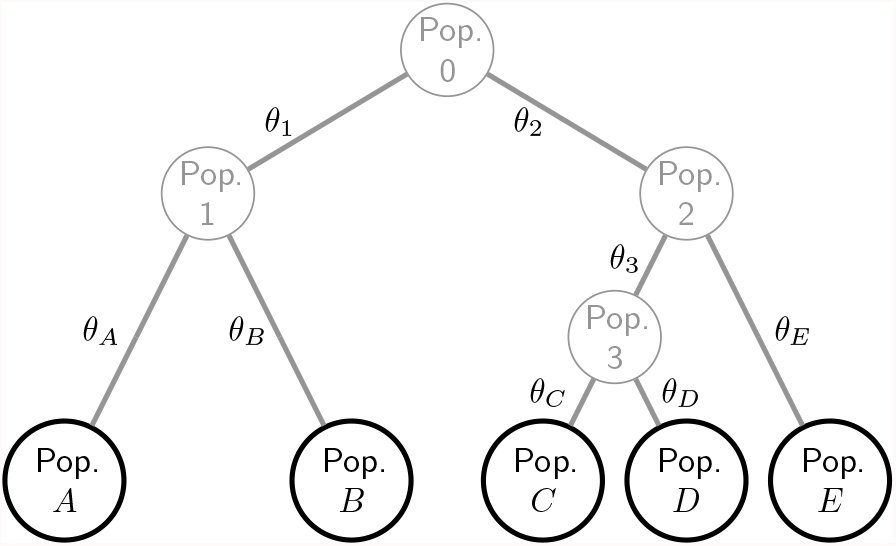
Hierarchically-structured populations. Global ancestral Population 0 and intermediate ancestral populations 1 to 3 are unobserved, while allele frequency data is available for leaf populations *A* to *E*. The label of a population also refers to the branch above that population.The genetic differences among the nine populations are described by eight *θ* values, one for each tree branch.

We continue to assume (1) in the population tree, with *f* replaced by the allele fraction in the parent population if this is not Population 0. See Table 1 for explanation of notation used in, and S1 Appendix for proofs of, the following results:

**Table 1.** Notations for population trees, with examples from the tree of Fig. 2.

| Definition | Examples from Fig. 2 |
| --- | --- |
| $\mathcal{P}(k)$ = tree path from 0 to $k$ | $\mathcal{P}(A) = \{1, A\}; \mathcal{P}(C) = \{2, 3, C\}; \mathcal{P}(E) = \{2, E\}$ |
| $\mathcal{Q}(kk') = \mathcal{P}(k) \cap \mathcal{P}(k')$ | $\mathcal{Q}(k) = \mathcal{P}(k); \mathcal{Q}(CE) = \mathcal{Q}(EC) = \{2\}$ |
| $\mathcal{R}(kk') = \mathcal{P}(k) \setminus \mathcal{P}(k')$ | $\mathcal{R}(k) = \emptyset; \mathcal{R}(CE) = \{3, C\}; \mathcal{R}(EC) = \{E\}$ |

#### Proposition 1.

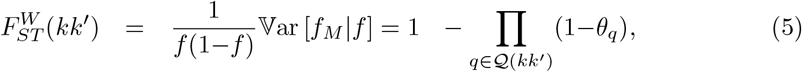

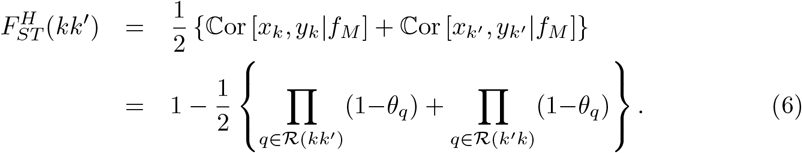

*where M denotes the most recent common ancestor of Populations k and k*^′^, *and a product over an empty set is defined to equal one*.

From Eqs (5) and (6) we see that 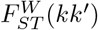 and 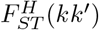 are functions of disjoint sets of *θ* coefficients. 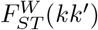 measures the shared genetic variation of populations *k* and *k*^′^ relative to Population 0, and so depends on *θ* values for tree branches between populations 0 and *M*. It can be interpreted as the probability of IBD from an ancestor between populations 0 and *M*. 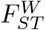 (*k*) reflects divergence of population *k* from the inferred ancestral population, while if *M* = 0 then 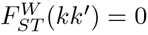..

Each product in (5) and (6) can be approximated by the inclusion-exclusion lower bound equal to one minus the sum of the *θ* values included in the product. This approximation is exact when the products include only one term.

Both 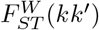 and 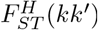 are invariant to switching *k* and *k*^′^. The value of 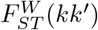, but not 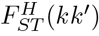, can change if new populations are included or existing populations (other than *k* and *k*^′^) are removed such that the ancestral population changes. The estimators 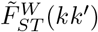 and 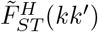, introduced below, depend on data from all the leaf populations, and both can be affected by addition or removal of populations other than *k* and *k*^′^.

#### The pairwise estimator of 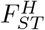

At a given locus, the pairwise estimator [5] is 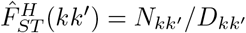, where

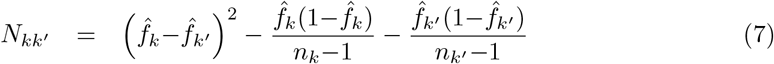

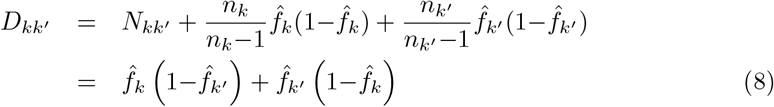

and *n*_*k*_ is the number of gametes sampled in population *k*, while 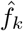 is the sample allele fraction. By expanding 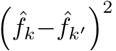, both *N*_*kk*′_ and *D*_*kk*′_ can be expressed as sums of terms of the form 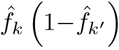 which for *k*≠*k*^′^ is an unbiased estimator of *f*_*k*_ (1−*f*_*k*′_). Assuming the conditional moments (1) we obtain

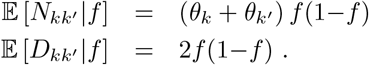

From (4), we see that the ratio of the above two expectations is 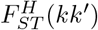 and so, provided that *D*_*kk*′_ has a low coefficient of variation which is typically the case in practice, 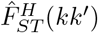 is approximately unbiased for 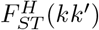.

*F*_*ST*_ can vary over SNVs, due to locus-specific effects of selection or mutation. In humans, there are relatively few strong outlier SNVs, and these can be removed prior to analysis if required, so that locus-specific selection and mutation effects are often ignored for genome-wide inferences of *F*_*ST*_. The multi-locus 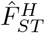 is defined by summing numerator and denominator over SNVs:

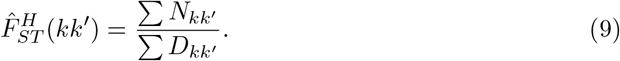

An unbiased alternative is to average the ratios over SNVs but, as previous authors have noted [2, 5, 9], the increased precision of the ratio of averages (9) more than offsets the small bias introduced. While our approach explicitly allows for linkage disequilibrium (LD) due to population structure, LD due to tight linkage is not modelled. Any effects of linkage on (9) are expected to be small in large, outbred species such as humans.

### Novel tree-based estimators

We construct a new method-of-moments estimator 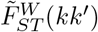 by first estimating the *θ* parameters using terms of the form 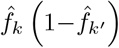. Let

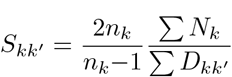

where 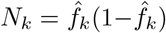 and *D*_*kk*′_ is defined at (8). The two summations are over the same set of SNVs, but any SNV that is monomorphic in *k* and *k*^′^ combined does not contribute to either sum and hence does not affect *S*_*kk*′_. While the fraction of monomorphic sites can be informative about *θ* values, data quality issues make it difficult to use this information in real datasets and it is ignored by our estimators.

#### Proposition 2.

*For k* ≠ *k*^′^,

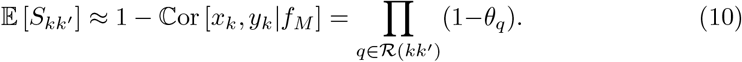

The result follows from the proof of Proposition (1): 𝔼 [*N*_*k*_] can be derived from the computation of ℙ[*x*_*k*_ ≠*y*_*k*_], and the derivation of 𝔼 [*D*_*kk*′_] corresponds to the computation of ℙ[*x*_*k*_ ≠ *x*_*k*′_].

Write *β*_*q*_ = log(1−*θ*_*q*_) and let *β* denote the vector of all *β*_*q*_ coefficients. We estimate *β* by solving:

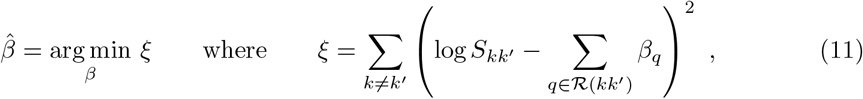

subject to *β*_*q*_ ≤ 0 since *θ*_*q*_ ≥ 0. For *K* populations, and noting that in general *S*_*kk*′_ ≠ *S*_*k*′*k*_, there are *K*(*K*−1) values of *S*_*kk*′_ available to estimate the 2(*K*−1) values of *β*_*q*_. Next we compute 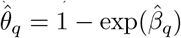 and define for all *k* and *k*^′^ (including *k* = *k*^′^)

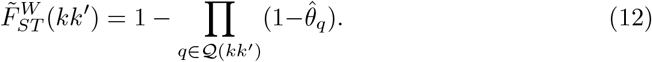

Finally 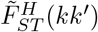 can be computed via (4) or (6).

### Fast inference of the population tree

We propose a fast algorithm to jointly infer the tree topology and *θ* values, which can be represented as branch lengths. Restricting the search to binary trees ensures that any two trees with *K* leaves have the same number of *θ* parameters to estimate, allowing the trees to be compared using the values of *ξ* in (11). A global search over all possible binary trees is infeasible even for moderate *K*. Instead, we first use a pairwise clustering strategy, starting with the independent-descent model (Fig. 1). At each step, an intermediate ancestral population is added between Population 0 and two of its child populations, chosen to minimise *ξ*. After *K*− 2 steps a binary tree is obtained.

Fig. 3 illustrates the clustering phase with *K* = 4 observed populations. We then seek to improve the tree obtained from the initial clustering. For each leaf node *k*, chosen in random order, we consider each branch in the current tree as an alternative location for the parent of *k*, fitting each of these 2(*K* −1) trees and choosing the one that minimises *ξ*, which may be the current tree in which case no change occurs.

**Figure 3.**
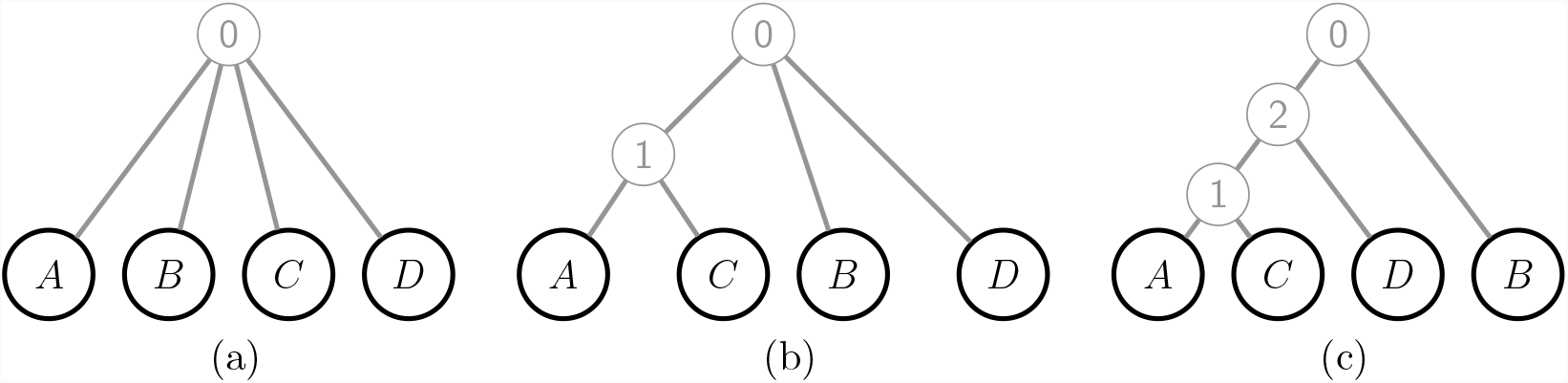
Hierarchical clustering to infer a binary tree for *K* = 4 observed populations. (a) The starting tree has all populations directly connected to ancestral Population 0. (b) We identify the pair of populations (here *AC*) such that an intermediate ancestral population between the pair and Population 0 minimises *ξ* in (11). (c) Repeating step (b), now 1*D* is the optimal pair of populations in {1, *B, D*}(the children of Population 0). After *K* − 2 = 2 steps, the resulting tree is binary and the algorithm stops.

In the clustering phase there are *K*−2 merge steps, and at the *j*th one there are 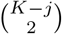 pairs of populations to consider merging. Overall we require 𝒪 (*K*^3^) solutions of the non-negative least-square optimization problem (11), for which we use the Lawson-Hanson algorithm [16]. The improvement phase of the algorithm scales with *K*^2^, because there are *K* leaves to consider relocating, and 2(*K* −1) locations to consider for each of them. In practice each step in the clustering phase can be solved easily using a warm-start strategy for initialization: each new fitting can be initialized using the tree and parameters inferred in the previous step. Consequently the actual computational burden of the improvement phase is usually higher. Solving (11) also requires computation of the *S*_*kk*′_, which is linear in *m*, the number of SNVs. This computation only has to be performed once, after which there is no further dependence on *m*, making the procedure feasible for any number of SNVs.

### Simulation study design

We randomly sampled gametes in leaf populations *A* to *E* that evolved according to the tree of Fig. 4. In the unobserved ancestral population (Population 0), the allele fraction *f* at an SNV was simulated from a beta(0.4,0.4) distribution, so that 19% of SNVs have MAF < 0.01 and 36% have MAF < 0.05. The allele fraction in each other population was simulated from a beta distribution with moments (1), using as *f* the allele fraction in its parent population.

**Figure 4.**
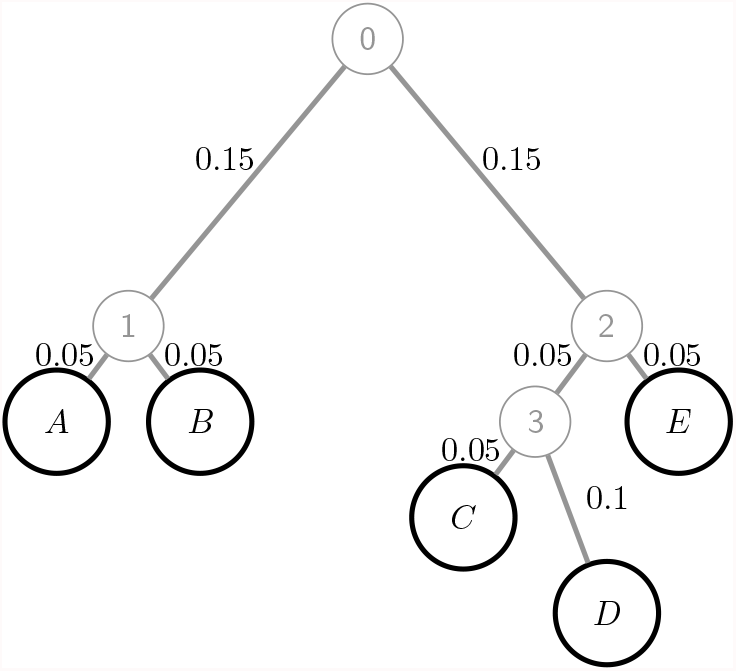
Population tree used for the simulation study. Each branch length *θ*_*q*_ is shown next to the corresponding branch.

The top two rows of Table 2 show simulation parameters, ordered from the most informative scenario (S1) to the least informative (S6) in terms of the number of correct tree inferences from 10 000 simulation replicates (Table 2, final row). While all population allele fractions remain positive under our model, genetic drift between Population 0 and the leaf populations increases the proportions of low-MAF SNVs, and many sites are monomorphic in the sample (Table 2, third row).

**Table 2.**
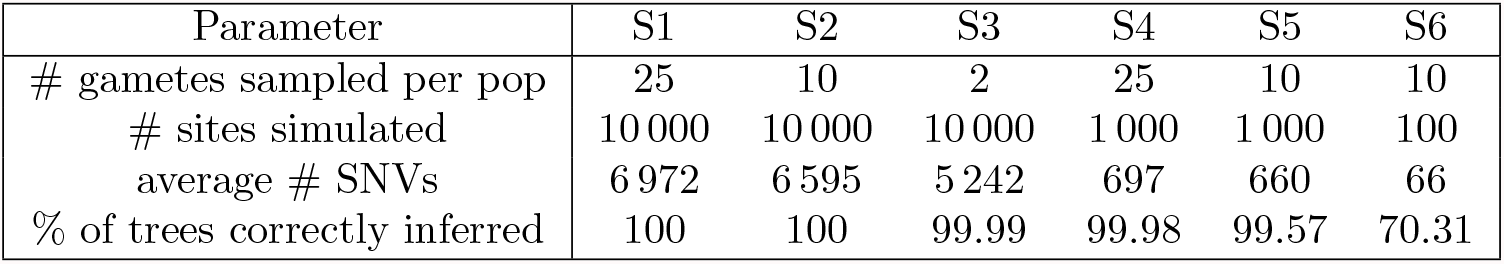
Details of six simulation scenarios based on the population tree of Fig. 4. # denotes “number of”. An SNV has at least one copy of a minor allele in at least one population, there is no MAF threshold.

Treating the tree structure in Fig. 4 as unknown, for each simulated dataset we jointly inferred the population tree and *θ* for each branch (see Methods), and then used (5) and (6) to compute our novel tree estimators 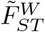 and 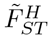, as well as the pairwise estimator 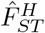 [5].

### The 1000 Genomes dataset

We applied our joint estimation of tree and *θ* values to data from phase 3 of the 1000 Genomes Project [17, 18] from 2 504 individuals sampled in 26 populations classified into five continental-scale “superpopulations” (Table 3). We included all available diallelic SNPs across the 22 autosomes, totalling 72M, with a binary coding indicating presence/absence of the major allele. The four AMR populations are strongly affected by historical admixture, including from different Native American source populations who are closest to the EAS superpopulation among the study populations. Estimated fractions of Native American ancestry are PEL 0.77, MXL 0.47, CLM 0.26 and PUR 0.13 [19]. The remaining ancestry comes mainly from European populations best represented among our study populations by IBS, but nearly 10% of the ancestry of AMR individuals is African (both European and African ancestry fractions are highest in PUR and lowest in PEL). ASW and ACB individuals also show some European admixture, but their ancestry is predominantly African (estimated fractions ACB 0.88, ASW 0.76 [19]). Some ASW individuals also show substantial Native American ancestry.

**Table 3.** Description of 1000 Genomes Project data [17, 18], and 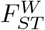 estimates. Next to each population code is a superpopulation label that is used for discussion but not in any analysis. 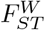 measures divergence of the population from the inferred global ancestral population.

| Population code | Description of ancestry; place sampled | Sample size (gametes) | $\tilde{F}_{ST}^W$ |
| --- | --- | --- | --- |
| ACB AFR | African Caribbeans; Barbados | 192 | 0.028 |
| ASW AFR | African Americans; SW USA | 122 | 0.054 |
| ESN AFR | Esan; Nigeria | 198 | 0.016 |
| GWD AFR | Gambian; Western Divisions, Gambia | 226 | 0.017 |
| LWK AFR | Luhya; Webuye, Kenya | 198 | 0.014 |
| MSL AFR | Mende; Sierra Leone | 170 | 0.015 |
| YRI AFR | Yoruba; Ibadan, Nigeria | 216 | 0.017 |
| CLM AMR | Colombians; Medellin, Colombia | 188 | 0.208 |
| MXL AMR | Mexicans; Los Angeles | 128 | 0.242 |
| PEL AMR | Peruvians; Lima, Peru | 170 | 0.303 |
| PUR AMR | Puerto Ricans; Puerto Rico | 208 | 0.191 |
| CDX EAS | Chinese Dai; Xishuangbanna, China | 186 | 0.307 |
| CHB EAS | Han Chinese; Beijing | 206 | 0.305 |
| CHS EAS | Southern Han Chinese | 210 | 0.308 |
| JPT EAS | Japanese; Tokyo | 208 | 0.307 |
| KHV EAS | Kinh; Ho Chi Minh City, Vietnam | 198 | 0.300 |
| CEU EUR | NW Europeans; Utah | 198 | 0.257 |
| FIN EUR | Finnish; Finland | 198 | 0.263 |
| GBR EUR | British; England and Scotland | 182 | 0.259 |
| IBS EUR | Iberians; Spain | 214 | 0.249 |
| TSI EUR | Toscani; Italia | 214 | 0.251 |
| BEB SAS | Bengali; Bangladesh | 172 | 0.231 |
| GIH SAS | Gujarati Indian; Houston, Texas | 206 | 0.237 |
| ITU SAS | Indian Telugu; UK | 204 | 0.234 |
| PJL SAS | Punjabi; Lahore, Pakistan | 192 | 0.230 |
| STU SAS | Sri Lankan Tamil; UK | 204 | 0.233 |

As well as the population-level analysis of all 2 504 individuals, we performed individual-level analyses for a subsample of five individuals from each of six populations: three AMR populations (CLM, MXL, PUR) and one population from each of AFR, EAS and EUR, namely the MSL, CHB and IBS populations. See S4 Table for identifiers of the selected individuals. Of the 72M SNVs in the full dataset, 13.4M remained SNVs in the 30-individual dataset. Of these 4.7M and 1.5M had one and two copies of the minor allele, respectively, while 1.7M had over 20 copies of the minor allele. We also performed principal component (PC) analysis which is a standard approach to visualising individuals based on their genome-wide genotypes. However, we did not apply the usual standardising of the SNV variables. Due to the absence of an MAF threshold, with standardisation the first five PCs are dominated by the 4.7M singleton sites and only differentiate the five MSL individuals from each other and the rest of the sample.

Computation for the 26-population and 30-individual analyses each took around 10 minutes. The different numbers of SNVs (72M and 13.4M) has little impact on computing time, and the first (clustering) phase of the tree-inference algorithm required just a few seconds for both analyses, with the improvement phase requiring most of the computing time.

## Results

### Simulation study

All the estimators considered here have low bias. The bias of 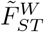 is undetectable for simulation S1, negligible for S3 and noticeable but small in the low-information S6 (Table 4). The RMSE of 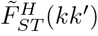 also tends to increase over the six scenarios (Table 5). In most scenarios it is lowest for the closest population pair *AB*, and highest for the two most divergent pairs, *AD* and *BD* (Table 5).

**Table 4.** 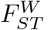 and estimated bias of our novel tree-based estimator 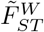 in three simulation scenarios. Based on 10 000 replicates in each simulation scenario (see Table 2). The values for *AD, AE, BC, BD*, and *BE* are the same as those for *AC*. When 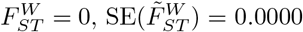 for S1 and S3, and = 0.0001 for S6. When 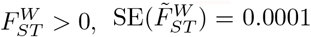 for S1 and S3, and = 0.0006 for S6.

| $kk'$ | $F_{ST}^W(kk')$ | Estimated bias of $\tilde{F}_{ST}^W(kk')$ | | |
| --- | --- | --- | --- | --- |
|  |  | S1 | S3 | S6 |
| $A$ | 0.192 | 0.000 | 0.000 | 0.001 |
| $AB$ | 0.150 | 0.000 | 0.000 | -0.005 |
| $AC$ | 0.000 | 0.000 | 0.000 | 0.001 |
| $B$ | 0.192 | 0.000 | 0.000 | 0.001 |
| $C$ | 0.233 | 0.000 | 0.000 | -0.001 |
| $CD$ | 0.192 | 0.000 | 0.001 | -0.006 |
| $CE$ | 0.150 | 0.000 | 0.000 | -0.002 |
| $D$ | 0.273 | 0.000 | 0.000 | -0.002 |
| $DE$ | 0.150 | 0.000 | 0.000 | -0.002 |
| $E$ | 0.192 | 0.000 | 0.000 | -0.007 |

**Table 5.**
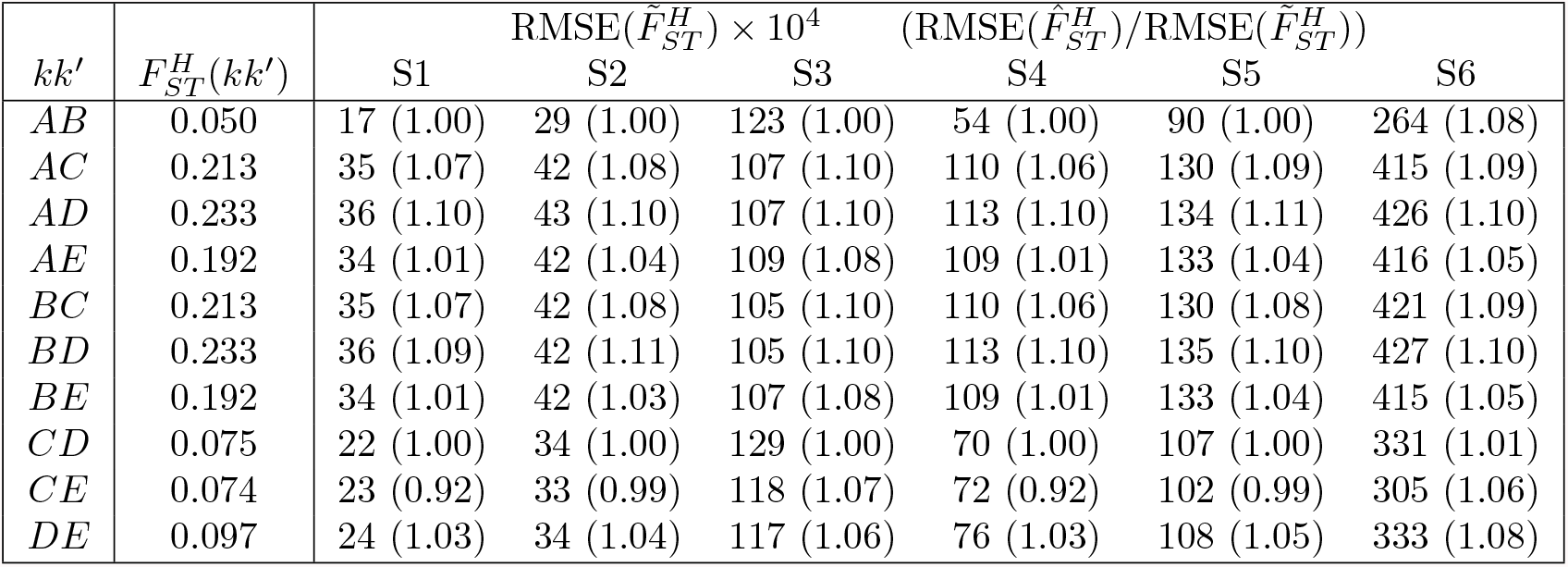
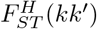 and the RMSE of 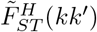. Based on 10 000 replicates of each of simulation scenario. In brackets is the ratio of the RMSE of 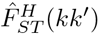, the pairwise estimator of [5], to that of 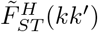; values *>* 1 indicate that 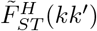 performs better than 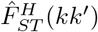.

Measured by RMSE, 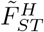 is superior to 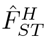 for 56 of the 60 values reported (Table 5). The superiority of 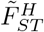 over 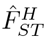 tends to be higher as the informativeness of the datasets declines. 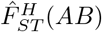 performs almost as well as 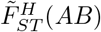, reflecting that there are no populations sufficiently close to *A* and *B* to provide useful information. Conversely, for all the population pairs only connected in the population tree via the root, 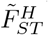 is superior to 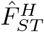. For example, in estimating 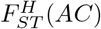, allele frequencies in population *B* are informative about frequencies in the path between *A* and *C*, and only 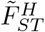 exploits this information. The exception is the pair *CE*, for which the allele frequencies in *D* convey some relevant information, but it does not always improve inferences of 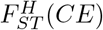, reflecting that *D* is highly diverged from the path connecting *C* with *E*.

In S3, the five observed populations are each represented by a sample of size two gametes, and so 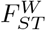 and 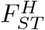 describe the coancestry among five individuals. For S6 with only 66 polymorphic SNVs, the population tree was correctly inferred in only 70% of simulations (Table 2, final row), but enough correct features of the tree were extracted to improve inference such that 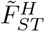 showed the largest relative improvement over 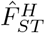 in this low-information scenario (Table 5).

### 1000 Genomes population analysis (26 populations, *n* = 2 504)

The single-population 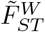 values, measuring the divergence of each of the 26 populations from the inferred global ancestral population (Table 3), are lowest for the AFR populations (0.01 – 0.05) and highest for PUR and the EAS populations (0.30 – 0.31). Greater divergence of non-AFR populations may reflect an out-of-Africa bottleneck. The AMR superpopulation has the greatest range of 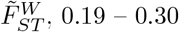, with values ordered by the level of African admixture. The average of the four values is 0.24, close to the average individual-specific *F*_*ST*_ of 0.23 for AMR reported by [20].

The 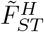 values, measuring divergence between pairs of leaf populations, show a familiar pattern for human population genetic studies, with the largest values comparing AFR with non-AFR populations, particularly the 35 AFR-EAS population pairs (Fig. 5, left). The largest value is 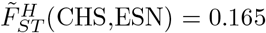. Within superpopulations, the maximum 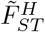 values are 0.007 for SAS, 0.011 for EUR, 0.013 for EAS, 0.031 for AFR and 0.068 for AMR.

**Figure 5.**
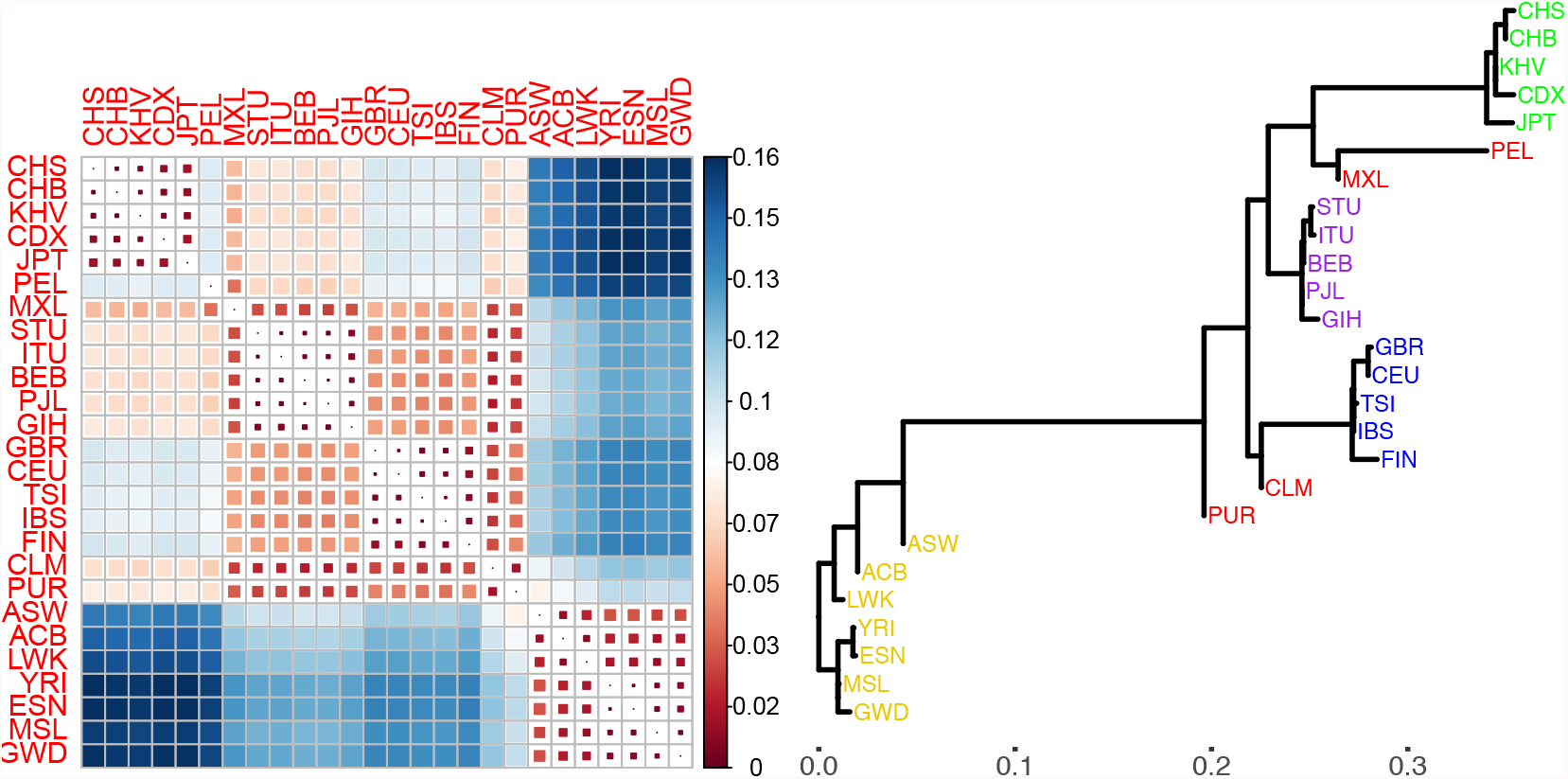
Tree-based inferences from the 26 populations of the 1000. Genomes dataset. Left: 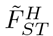 values (see scale for colour coding) for each pair of populations. Right: The inferred population tree, with horizontal branch lengths corresponding to coancestry parameter estimates 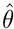 (see *x*-axis scale). Vertical distances have no meaning and are for display purposes only.

The inferred population tree (Fig. 5, right) reveals more structure than is evident from the matrix of 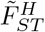 values. As expected, the AFR, EAS, EUR and SAS superpopulations each cluster together, but now we can also see that the admixed AFR populations, ACB and ASW, are closer to non-AFR populations, and IBS is the EUR population closest to non-EUR populations, reflecting the contribution of Iberians to AMR populations. The longest branch in the tree, which connects ASW with PUR, lies on the path between every AFR and non-AFR population pair, giving PUR a central position among the 1000 Genomes populations. Consequently, 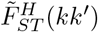 where *k* ∈ AFR and *k*^′^ ∉ AFR is well approximated by 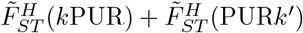.

The root of the inferred tree separates West African (ESN, GWD, MSL, YRI) from all other populations, consistent with the origins of modern humans in Africa. The largest genetic distances are between West-African and EAS populations, which reflects their geographical separation and low historical migration.

Fig. 5 (right) gives a visual representation of actual genetic variation among the 1000 Genomes populations. The observed variation is strongly influenced by historical processes of splitting and divergence, but the tree may not accurately reflect actual historical events. For example, the inferred global ancestral population provides a convenient reference for describing components of genetic variance, but may not accurately represent any actual population in human history.

While admixture events are not explicitly modelled, effects of admixture can be discerned in the current patterns of genetic similarity. The AMR populations are divergent from each other and other populations, with CLM closest to EUR and PEL and MXL closest to EAS populations, corresponding with their levels of admixture outlined above. PEL is the population most distant from its nearest neighbour, MXL, with 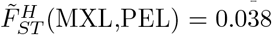.

The high correlation of 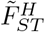 and 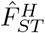 values (0.984) is driven by the similarity of the two estimators for the largest genetic distances (Fig. 6, left), whereas there are substantial differences between them over most of the range. The comparisons between PUR and the five EUR populations give the largest values of 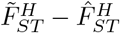, between 0.027 and 0.030 (Fig. 6, right). There are three comparisons for which 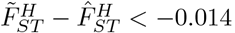, each involving a EUR and an EAS population (TSI-CDX, KHV-IBS, and CDX-IBS).

**Figure 6.**
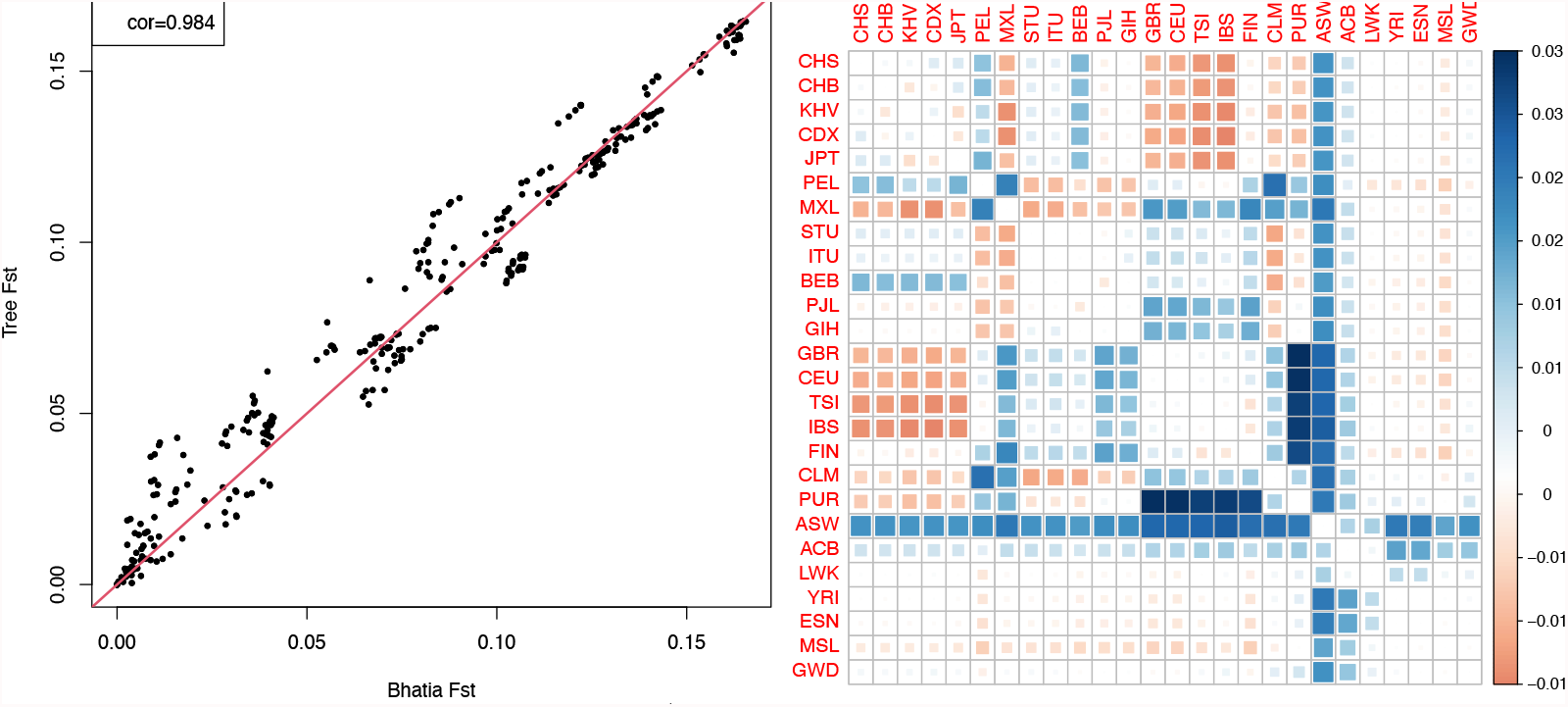
Left: pairwise estimator 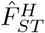 [5] (x axis) and tree-based estimator 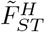 (y axis) for all 325 pairs of populations in the 1000 Genomes dataset. Right: colour-coded values of 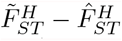, which is largest for comparisons of PUR with all 5 EUR populations (dark-blue squares).

### 1000 Genomes individual analysis (6 populations, *n* = 30)

There is good agreement in 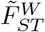 values between the individual and population analyses (Table 6), despite the great difference in sample size and populations sampled. This is important because 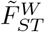 is based on the inferred ancestral population, but estimates can still be comparable across very different observed datasets.

**Table 6.**
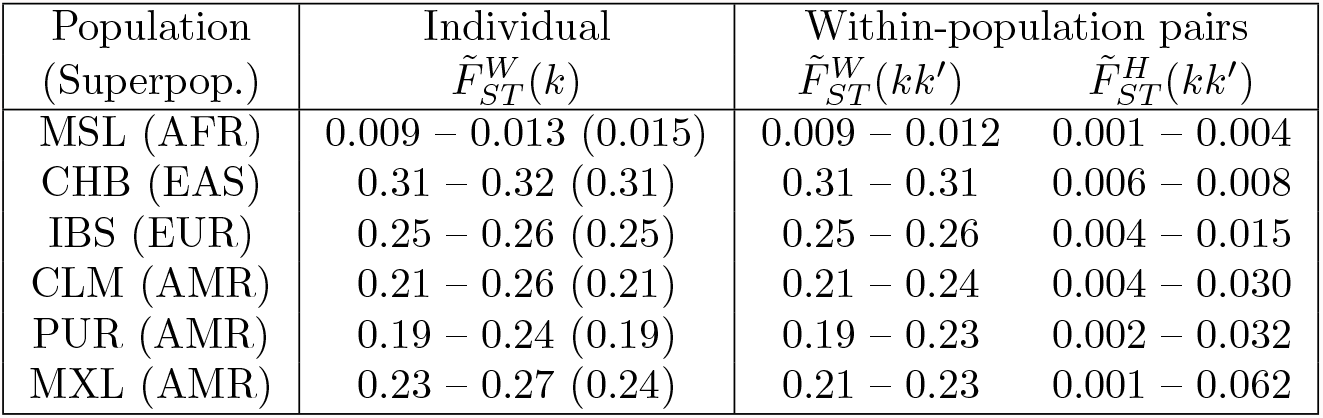
Column 2 gives the range over 5 individuals from the population in column 1 of 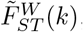, measuring divergence from the inferred ancestral population, and (in brackets) the corresponding population-level value from Table 3. Also shown are the ranges over the 10 within-population pairs of individuals of 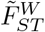 and 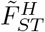, measuring respectively shared genetic variance and between-pair divergence.

Fig. 7 shows a PC plot and the inferred tree for the 30 individuals. The two plots convey similar information, with the tree giving finer detail about shared and non-shared components of variance among the individuals plus interpretability from horizontal branch lengths corresponding to *θ* and hence *F*_*ST*_ estimates. The CLM, MXL and PUR population labels indicate location of sampling, but they do not accurately reflect genetic structure because of the high within-group diversity, with many instances of between-group pairs of individuals being genetically closer to each other than within-group pairs. One MXL individual is genetically closer to all of the IBS sample than to any other MXL individual.

**Figure 7.**
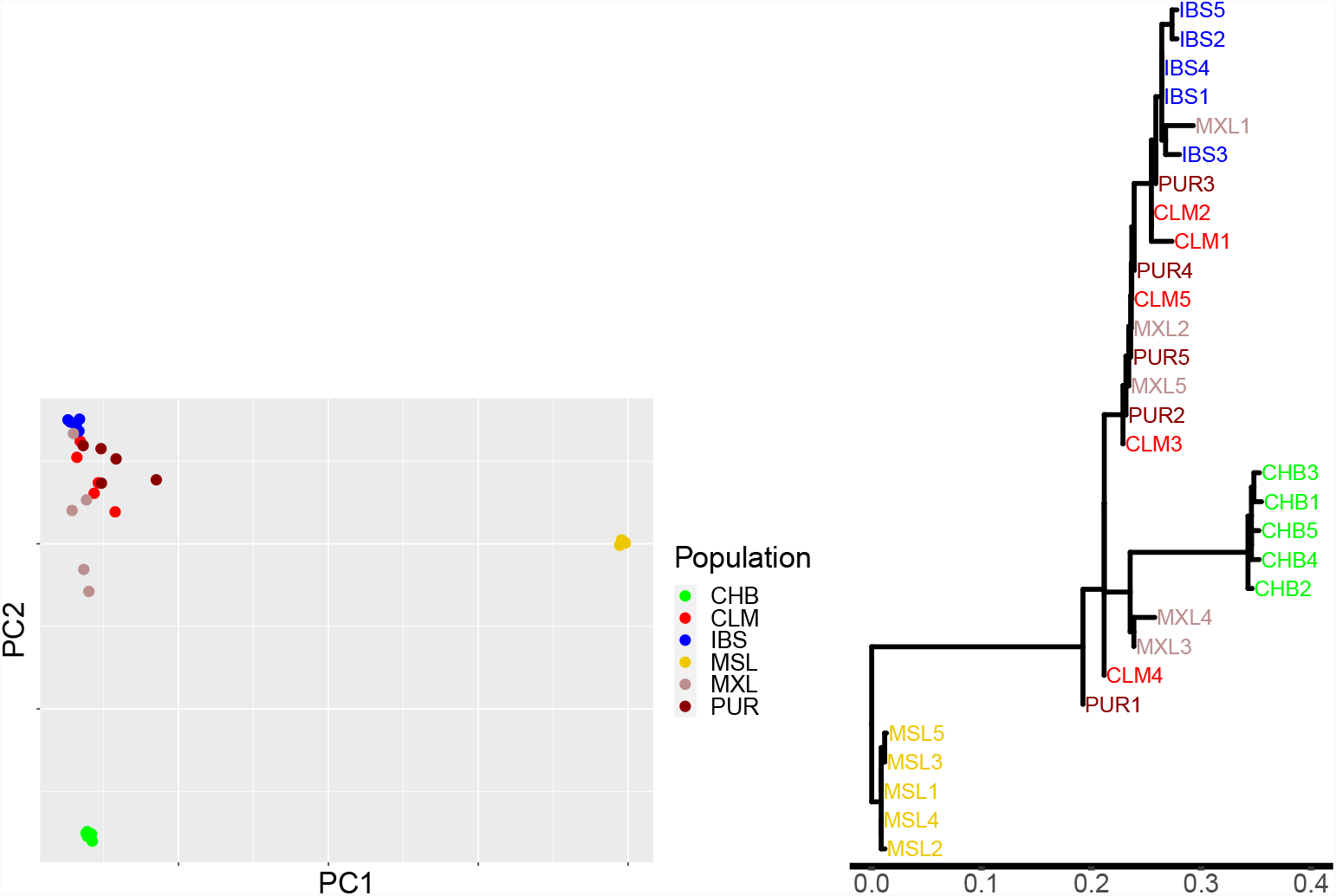
Left: First two principal components (explaining 29% of variance) from 13.4M unstandardised SNVs in a sample of 30 individuals from the 1000 Genomes dataset (5 each from six populations as indicated in the legend box). Right: inferred tree for the 30 individuals, with horizontal branch lengths corresponding to coancestry parameter estimates 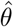.

Table 6 shows a wide range of between-individual divergence 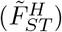 in the admixed AMR populations. The non-AMR populations are more homogeneous, although there is higher between-pair divergence in IBS than in CHB or MSL, which may reflect some migration from the Americas. S3 Fig shows the corresponding tree when the non-AMR individuals are pooled into 3 population samples. This analysis reduces computing time with little loss of information.

## Discussion

We have extended the definitions of the genetic distance *F*_*ST*_ to the tree-structured multi-population setting, showing that correlation and mismatch probability definitions of *F*_*ST*_ measure shared and non-shared genetic variation since the most recent population ancestral to all observed populations. We then developed fast and efficient estimators of these parameters, and an algorithm to infer the topology of a tree of ancestral populations that models the observed population structure. Although methods for inferring a population tree from allele frequency data are already available, including `Treemix` [21] and Neighbor Joining [22], our procedure is the first to perform joint inference of *F*_*ST*_ and the tree, which allows sharing of information about allele frequencies in ancestral populations and a range of options for visualising genetic structure by combining homogeneous groups of individuals.

The inferred tree will typically reflect the large-scale evolutionary processes of population splitting and divergence, but it may not accurately reconstruct the evolutionary history of the populations studied because there is no explicit role for migration following a population split. Instead, the tree provides a visual representation of the actual genetic variation across the populations, with the added advantage of branch lengths that are interpretable as *θ* values that specify the contribution from that branch to *F*_*ST*_ values. Locus-specific *F*_*ST*_ values that diverge from genome-wide averages have long been used to help identify the effects of natural selection [23–26], with some methods depending explicitly on a population tree [27, 28]. Our improved tree inference and parameter estimates may be able to increase the power of such methods, and simultaneous inference of all *θ* parameters should lead to better characterisation of the selection effect, which will be explored in future research.

Our methods also provide a novel approach to describing coancestry among sets of diploid individuals, treating each as a population of two gametes. This is not practical for pairwise estimators of *F*_*ST*_ because of inadequate information about reference allele frequencies, whereas other individuals and populations inform about them in our approach. Any pair of individuals are related through many ancestral lineages of varying lengths. Pedigree-relatedness captures only very short lineage paths (within the known pedigree), whereas 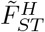 for two individuals is affected by all lineage paths connecting them, which can be useful to construct adjustments for even subtle population structure in heritability analyses and genetic association analyses. Currently we do not model LD among the markers and so cannot accommodate closely related individuals, but close relatives are also usually excluded from genome-wide association studies.

Since the seminal contribution of Lewontin [29], there has been interest in comparing genetic diversity within and between human populations. For example, within-Africa genetic differences were reported to be larger than differences between Eurasians and Africans [30]. Estimates 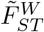 provide a convenient way to quantify such comparisons. Figure 7 (right) shows that, for these six populations, diversity is much lower within CHB and MSL than for any between-population comparison, but for CLM, MXL and PUR within-population and between-population 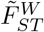 can be of similar magnitude.

To simplify the presentation we have not here inferred inbreeding parameters, but we expect this to be a straightforward extension. Another extension is to model the component of variance shared by a set of populations or individuals, rather than just pairs, which requires replacing 𝒬 in (5) with 𝒫 (*k*_1_) ∩ 𝒫 (*k*_2_) ∩ … ∩ 𝒫 (*k*_*p*_).

Values of 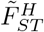 for pairs of individuals, illustrated graphically in Fig. 7 (right), can be useful in assessing forensic match probabilities comparing alleged and alternative sources of a crime-related DNA sample [31]. For practical reasons *F*_*ST*_ has been estimated at the level of populations [32, 33] but the relevant value of *F*_*ST*_ measures relatedness of pairs of individuals, the alleged and true sources of a crime-related DNA sample. While it is not typically possible to estimate *F*_*ST*_ for this specific pair of individuals, since the identity of the true source is a matter of dispute, a range of *F*_*ST*_ values over many pairs of individuals can indicate values that may be relevant to a particular case. We can also include population data from a forensic database, which can be used to ensure a representative reference population that is the same over different cases. Forensic DNA profiling primarily uses short tandem repeat loci rather than SNVs, and these have different 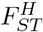 values due to a different mutation process, but SNV-based DNA profiling is becoming more common [34].

## Supporting information

Statistical analyses were performed using R. The main functions to compute Fst estimates and perform tree inference can be found in the R package HFst, available on the personal webpage of the first author https://www6.inrae.fr/mia-paris/Equipes/Membres/Tristan-Mary-Huard along with a short tutorial to perform the population analysis based on chromosomes 21 and 22.

## S1 Appendix. Proof of Proposition 1

### Lemma 1.

*Let F*_1_, …, *F*_*K*_ *be random variables satisfying for k* ∈ [1 : *K*−1]:

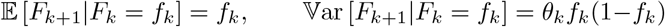

*Then*

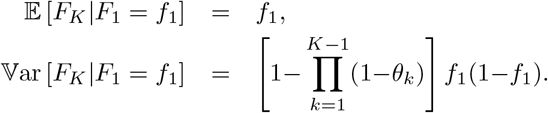

The proof is by induction. The case *k* = 1 is given as an assumption in the statement of the Lemma. Assuming that the required property is satisfied for *k* = *K*−1, and writing *g* for the conditional probability density function of the *F*_*k*_, we can express 𝕍ar [*F*_*K*_|*F*_1_=*f*_1_] as:

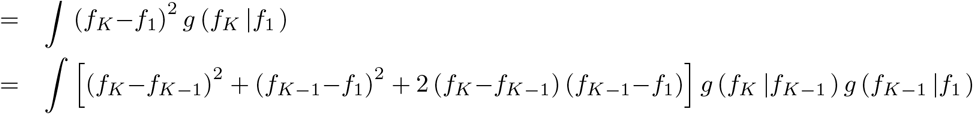

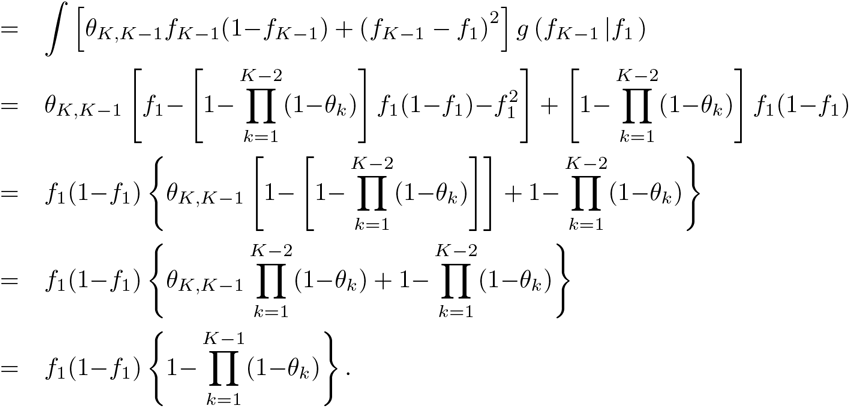

Now recall the definition of 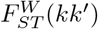 in (2), which can be expressed as:

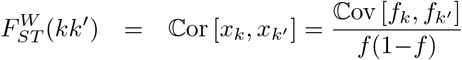

and, recalling that *M* denotes the most recent common ancestor of *k* and *k*^′^,

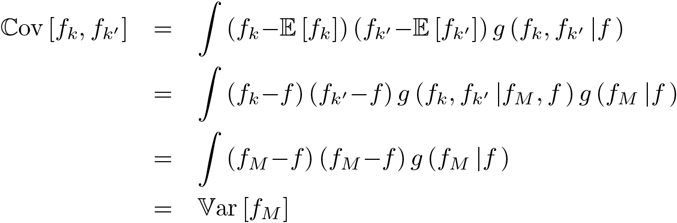

which establishes the first equality, then Lemma 1 provides (5).

Now recall the definition of 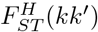 at (3). Focusing first on the probabilities in the denominator,

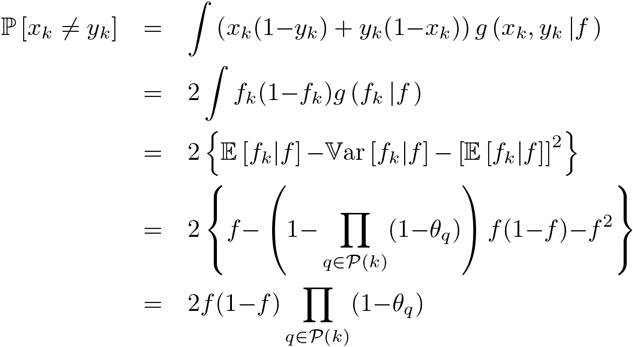

where the last equality comes from Lemma 1. A similar computation for the denominator leads to

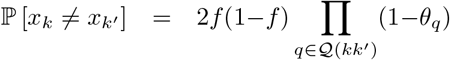

Therefore,

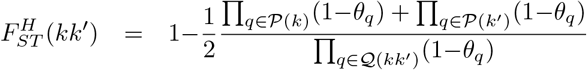

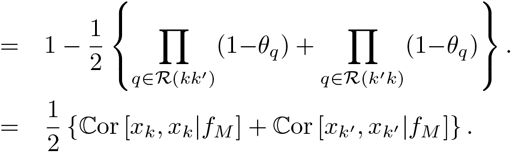

## S2 Appendix. Details of the tree-based inference algorithm

Here we use the notations of Table 1, now including subscripts such as *T* to identify the current tree. A tree-like population model *T* is characterized by the set of paths to each leaf population from ancestral Population 0,{𝒫_*T*_ (*k*), *k* ={*A*, …, *K*}}, together with *θ* values for each tree branch.

### Phase 1: Clustering

We propose an ascending clustering strategy to obtain an approximately optimal 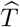. We introduce

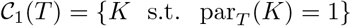

the set of all (observed or ancestral) populations whose parent population is ancestral Population 0. The ascending algorithm, also illustrated on a 4 population example in Fig. 3, works as follows:

Step 1 *Initialization*: Set *j* = 1 and let *T*_1_ denote the independent-descent population model (Fig. 1), so that

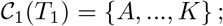

Step 2 *Merging* : Increment *j*. Consider each pair {*c*_1_, *c*_2_} ∈ 𝒞_1_(*T*_*j*−1_) and add an intermediate ancestral population *j* in tree *T*_*j*−1_ to create a new tree 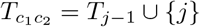 such that 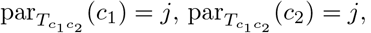, and 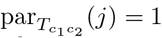. Infer the *β* for each 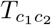 using (11), and then set 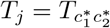 where

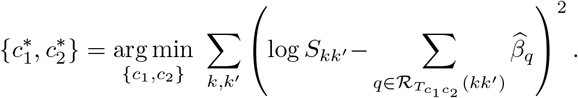

Step 3 *Stopping* : If *j* = *K*−1 set 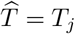 and stop, else go to step 2.

### Phase 2: Improvement

We attempt to update 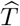 by choosing a leaf node *k* in random order, and consider relocating its parent to each branch of the current tree, choosing the branch that minimizes *ξ*.

**S3 Fig.**
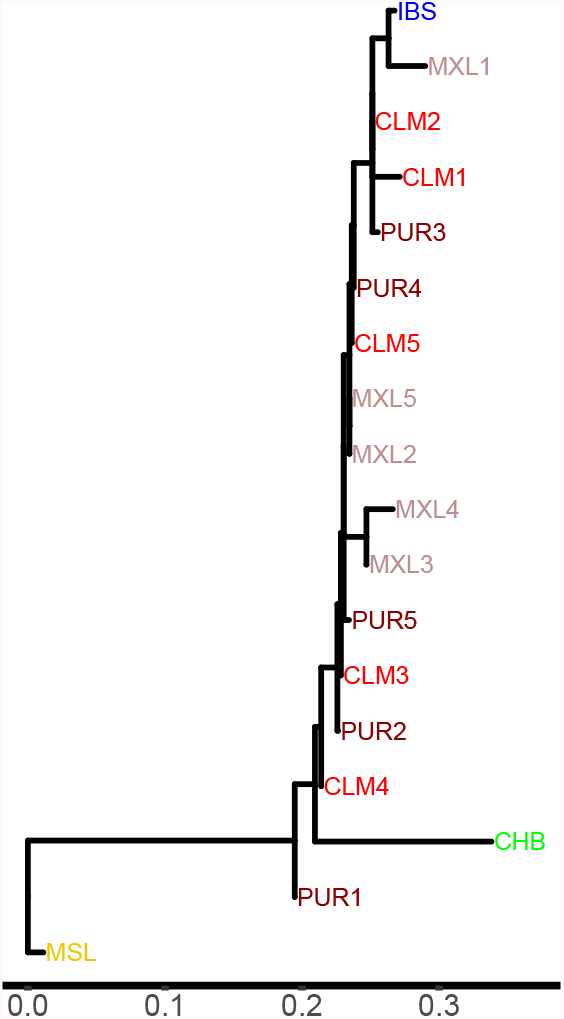
Inferred tree showing coancestry among the 15 individuals and 3 population samples. Similar to Fig. 7 (right) except that the 5 individuals from CHB, IBS and MSL samples have been pooled.

**S4 Table.** 1000 Genomes Project identifiers for the 5 individuals from each of 6 populations in the individual analyses.

| Population | Individual ID |
| --- | --- |
| CLM | HG01112 HG01119 HG01122 HG01125 HG01131 |
| MXL | NA19648 NA19652 NA19661 NA19681 NA19684 |
| PUR | HG01161 HG01164 HG01168 HG01173 HG01177 |
| CHB | NA18525 NA18528 NA18531 NA18533 NA18535 |
| IBS | HG01500 HG01503 HG01506 HG01509 HG01512 |
| MSL | HG03052 HG03058 HG03063 HG03069 HG03079 |

## Acknowledgments

This research was partially supported by grant DP190103188 from the Australian Research Council to DB, and by the “Investissement d’ Avenir” project (Amaizing, ANR-10-BTBR-0001) to TMH.

The authors gratefully acknowledge Dr Angad Johar for assistance with extracting the 30-individuals subset of the 1000 Genomes Project data.

